# Predicting microbial genome-scale metabolic networks directly from 16S rRNA gene sequences

**DOI:** 10.1101/2024.01.26.576649

**Authors:** Ove Øyås, Carl M. Kobel, Jon Olav Vik, Phillip B. Pope

## Abstract

Genome-scale metabolic models are key biotechnology tools that can predict metabolic capabilities and growth for an organism. In particular, these models have become indispensable for metabolic analysis of microbial species and communities such as the gut microbiomes of humans and other animals. Accurate microbial models can be built automatically from genomes, but many microbes have only been observed through sequencing of marker genes such as 16S rRNA and thus remain inaccessible to genome-scale modeling. To extend the scope of genome-scale metabolic models to microbes that lack genomic information, we trained an artificial neural network to build microbial models from numeric representations of 16S rRNA gene sequences. Specifically, we built models and extracted 16S rRNA gene sequences from more than 15,000 reference and representative microbial genomes, computed multiple sequence alignments and large language model embeddings for the 16S rRNA gene sequences, and trained the neural network to predict metabolic reaction probabilities from sequences, alignments, or embeddings. Training was fast on a single graphics processing unit and trained networks predicted reaction probabilities accurately for unseen archaeal and bacterial sequences and species. This makes it possible to reconstruct microbial genome-scale metabolic networks from any 16S rRNA gene sequence and enables simulation of metabolism and growth for all observed microbial life.

## Introduction

Genome-scale metabolic models (GEMs) allow simulation of metabolism and growth, with a wide range of applications from microbial species and communities to multi-cellular eukaryotes [1]. Reconstructing eukaryotic metabolism still requires substantial manual curation [2], but automated reconstruction of accurate GEMs from genomes is possible for microbes [3, 4]. Fueled by rapidly growing genomic and biochemical databases [5, 6], automated metabolic reconstruction methods have produced large GEM collections, primarily of human gut microbes [7–9] but recently of unicellular fungi as well [10]. However, despite progress in metagenomics [11] and cultivation [12], many observed microbes lack genomic data and thus remain inaccessible to GEMs.

Marker genes such as 16S ribosomal RNA (rRNA) allow estimation of relative microbial abundances through operational taxonomical units (OTUs) or amplicon sequence variants (ASVs), and they are commonly used for taxonomic and functional profiling of microbiomes [13]. Metagenome-assembled genomes (MAGs) improve microbiome analyses but technical and financial barriers can limit their ability to assess taxonomy and function for all microbes in a sample [14]. This also hampers functional prediction from 16S rRNA sequencing data, which currently relies on taxonomic mapping of OTUs or ASVs to genomes for predicting functional category abundances [15, 16] or to GEMs built from these genomes for predicting metabolic reaction abundances [17–19]. The fact that many observed microbes are unamenable to functional analyses necessarily adds bias towards well-studied and cultured microbial taxa.

Here, we leveraged automated metabolic reconstruction methods and more than 15,000 high-quality microbial genomes to develop a machine learning approach that predicts microbial GEMs directly from 16S rRNA gene sequences. Specifically, we trained an artificial neural network (ANN) on pairs of GEMs and 16S rRNA gene sequences, multiple sequence alignments (MSAs), or large language model (LLM) embeddings (**Fig. 1**). The trained ANNs predicted GEMs accurately from archaeal and bacterial 16S rRNA gene sequences, showing that genome-scale metabolism can be reconstructed from the sequence of an individual gene and thus extending the scope of GEMs to all microbes observed through marker genes.

**Fig. 1.**
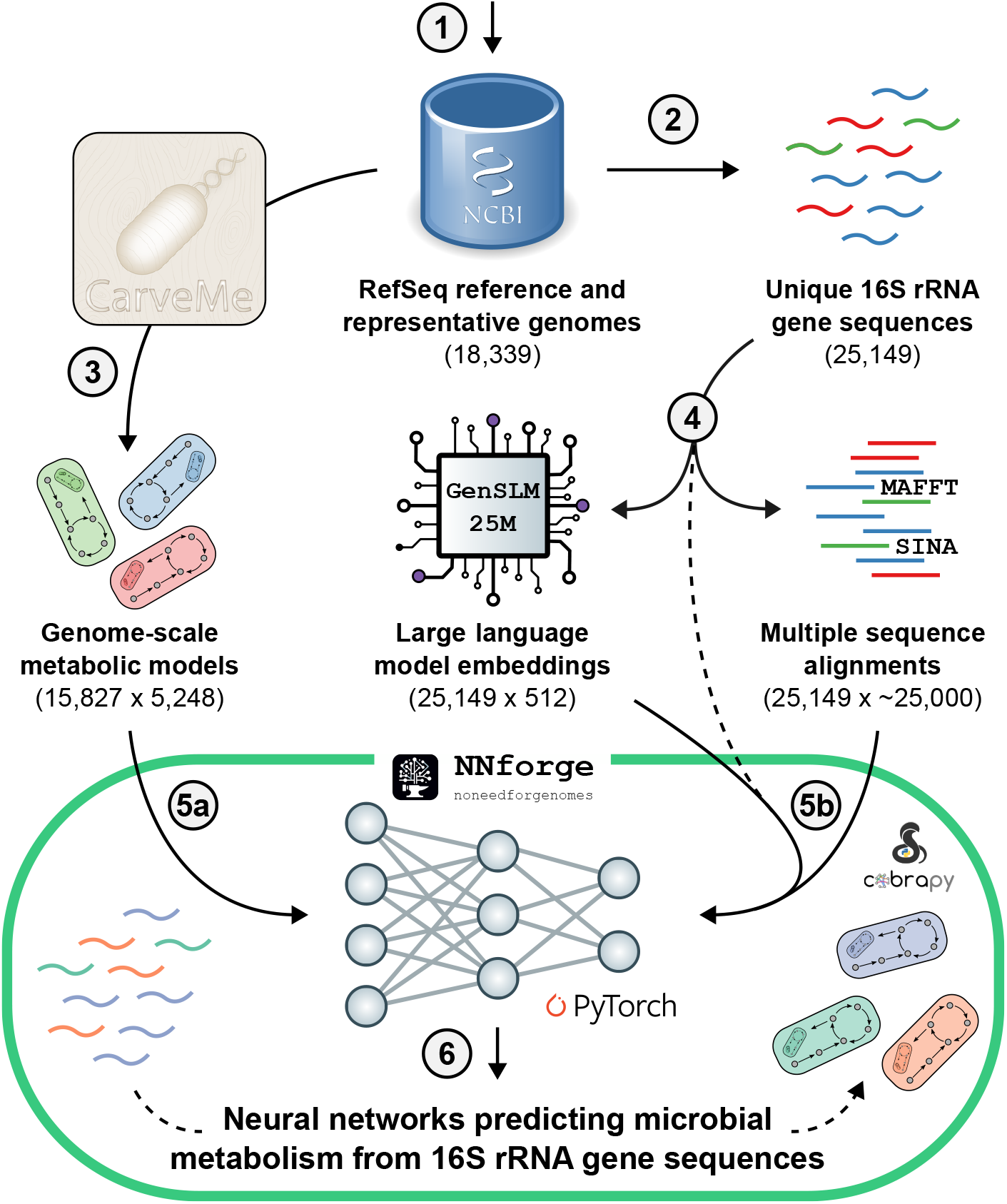
Predicting microbial GEMs from 16S rRNA gene sequences representations. We (1) downloaded all reference and representative genomes from RefSeq, (2) extracted all 16S rRNA gene sequences, and (3) built GEMs from the genomes using CarveMe. Then, we (4) represented the 16S rRNA gene sequences numerically as unaligned sequences, LLM embeddings, and MSAs from the aligner SINA and MAFFT, before training an ANN on pairs of (5a) GEMs and (5b) 16S rRNA gene sequence representations. The trained ANNs can be used to (6) predict microbial GEMs directly from 16S rRNA gene sequence representations.

## Results

### GEMs for reference and representative prokaryotic genomes

We downloaded all 18,339 available reference and representative genomes of archaea and bacteria from RefSeq [5], 15,827 of which could be mapped to at least one 16S rRNA gene sequence. The genomes covered seven of the 21 archaeal phyla (33%) in the Genome Taxonomy Database (GTDB) [20] and 52 of the 181 bacterial phyla (29%), and this coverage decreased with taxonomic rank to 12% of archaeal species and 18% of bacterial species (**Fig. 2**). The genomes covered the major clades of both archaea and bacteria, but some were covered by thousands of genomes and others only by a few. We built GEMs capable of simulating metabolic fluxes and growth from the genomes using the automated metabolic reconstruction tool CarveMe [3].

**Fig. 2.**
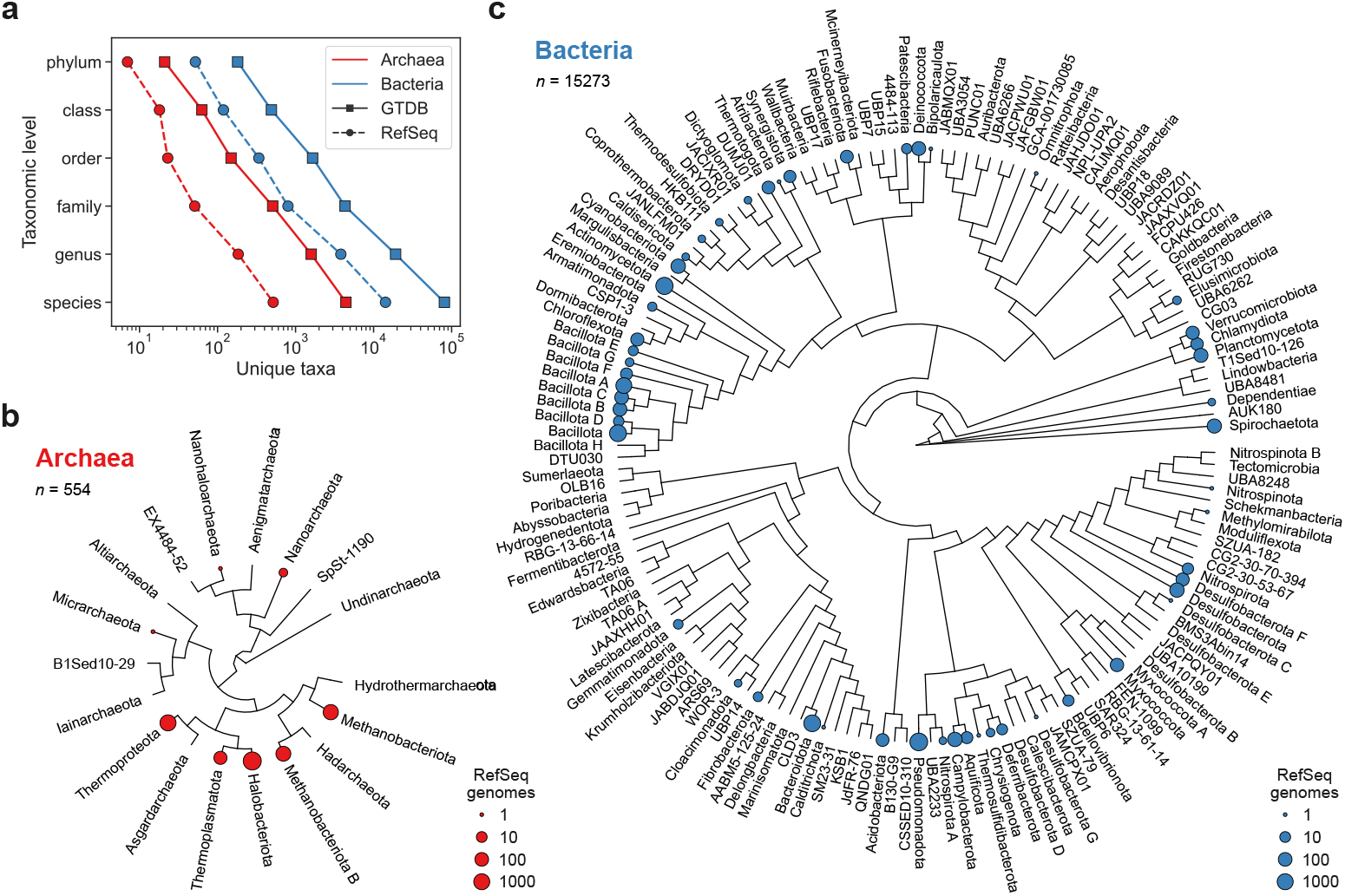
GTDB taxonomy coverage of reference and representative RefSeq genomes. (a) Number of taxa across taxonomic ranks for archaea and bacteria in the GTDB and RefSeq databases. The RefSeq genomes were filtered to only include reference and representative genomes that could be mapped to 16S rRNA gene sequences. (b) Cladogram of the GTDB reference tree for archaeal phyla with number of RefSeq genomes indicated per phylum. (c) Cladogram of the GTDB reference tree for bacterial phyla with number of RefSeq genomes indicated per phylum.

### Training an ANN on GEMs and 16S rRNA gene sequences

We developed a machine learning approach to predict GEMs directly from 16S rRNA gene sequences. From the annotation of the reference and representative RefSeq genomes, we extracted the sequences of 26,893 16S rRNA genes, which were represented numerically as (1) unaligned sequences, (2) LLM embeddings from GenSLM 25M [21], and (3) MSAs from the aligners SINA [22], which is specialized for 16S rRNA gene sequences, and MAFFT [23], which works for general sequences. Each GEM was represented by presence or absence of 5,248 unique reactions and was paired with 16S rRNA gene sequences from the same genome. We designed an ANN that gradually reduces the number of features over three fully connected linear layers, i.e., layers that connect every input neuron to every output neuron, with a final sigmoid layer that returns reaction probabilities. We trained this network on pairs of GEMs and 16S rRNA gene sequence representations. Specifically, we split our data into a training set (90%) and a test set (10%) ten times, trained the ANN on the training sets using each sequence representation, and used trained ANNs to predict for the test sets (**Fig. 3**). The network learned to predict GEMs accurately from all 16S rRNA gene sequence representations, but we did find significant differences between representations when predicting for the test set. Unaligned sequences produced the highest mean test loss, i.e., the worst reaction probability predictions across test sets, while LLM embeddings performed significantly better. MSAs gave the highest predictive accuracy, significantly better than LLM embeddings, but we found no significant difference between the two aligners, SINA and MAFFT. Predicted reaction probabilities for models in the test sets tended to be very close to zero for reactions that were absent in the true GEMs built from genomes and very close to one for reactions that were present. However, the distributions of predicted reaction probabilities had long tails, especially for reactions that were present in the true model. This was similar across all sequence representations, and predictions were indeed strongly and positively correlated between representations. On average, across all test sets, we found a Pearson correlation between sequence representations of 0.83 for reactions and 0.93 for GEMs, with strongest correlation between SINA and MAFFT.

**Fig. 3.**
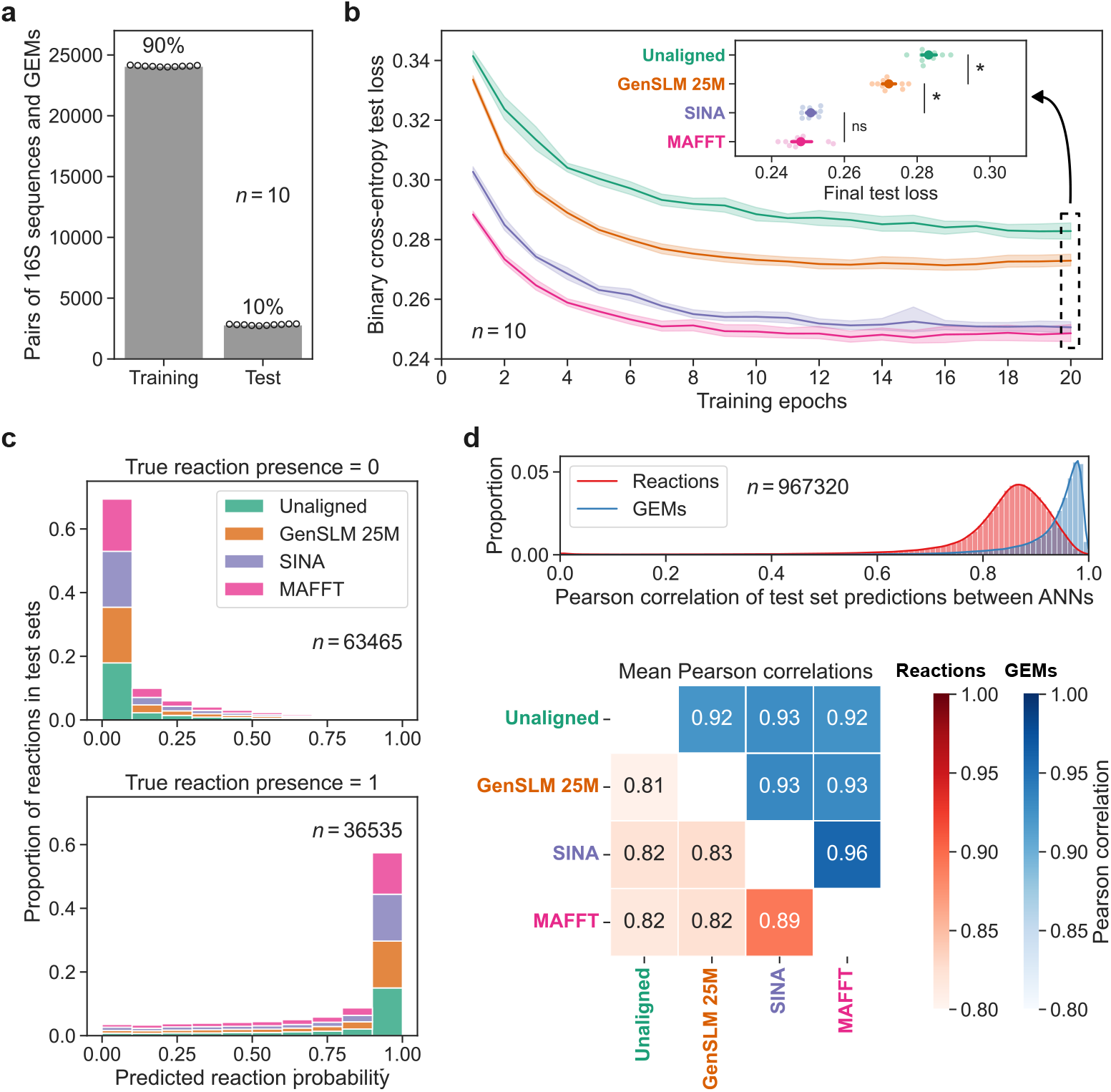
Training ANNs to predict GEMs from 16S rRNA gene sequence representations. (a) Sizes of sampled training and test sets. (b) Accuracy of ANN predictions for test sets by number of training epochs for different 16S rRNA gene sequence representations. The inset shows the final test loss with stars indicating significant differences (*p <* 0.05, paired *t*-test and Benjamini-Hochberg adjustment). Mean test loss is shown with 95% confidence interval of the mean from bootstrapping with 1,000 samples. (c) Predicted reaction probabilities for reactions that were absent (above) and reactions that were present (below) in models in the test sets. We randomly sampled 100,000 reactions from models in the test sets. (d) Distributions and means of Pearson correlations between ANNs trained on different 16S rRNA gene sequence representations for reactions (red) and GEMs (blue).

### Predicting GEMs accurately across archaea and bacteria

Converting reaction probability predictions into computable GEMs, i.e., models that can be solved by standard methods such as flux balance analysis [24], requires binary classification of reaction presence and absence. To do this, we systematically tested classification thresholds for all 16S rRNA gene sequence representations and test sets. For each GEM, we first classified reactions as present or absent using probability thresholds from zero to one. Then, we counted the number of true and false positives and negatives by comparing the classification to the true GEM built from the genome. Finally, we computed the area under the curve (AUC) of the receiver operator characteristic (ROC) curve as well as the Matthews correlation coefficient (MCC).

Trained ANNs predicted GEMs accurately and robustly across archaeal and bacterial phyla (**Fig. 4**). For ANNs trained on MSAs from SINA and MAFFT, the AUC of the ROC curve for classification of GEMs in the test sets was 0.94 for archaea and 0.96 for bacteria, meaning that there were very few false positives compared to true positives across classification thresholds. The threshold *α* = 0.5, which approximately maximized the MCCs of GEMs in the test sets, produced a median MCC of 0.71 for archaea and 0.76 for bacteria. Most predicted GEMs were accurate, although the MCC distributions had long tails of less accurate GEMs, most notably for archaea. Predictions were consistently accurate for all but a few archaeal and bacterial phyla: median MCC was greater than 0.7 for 33% of the phyla and greater than 0.6 for 70%.

**Fig. 4.**
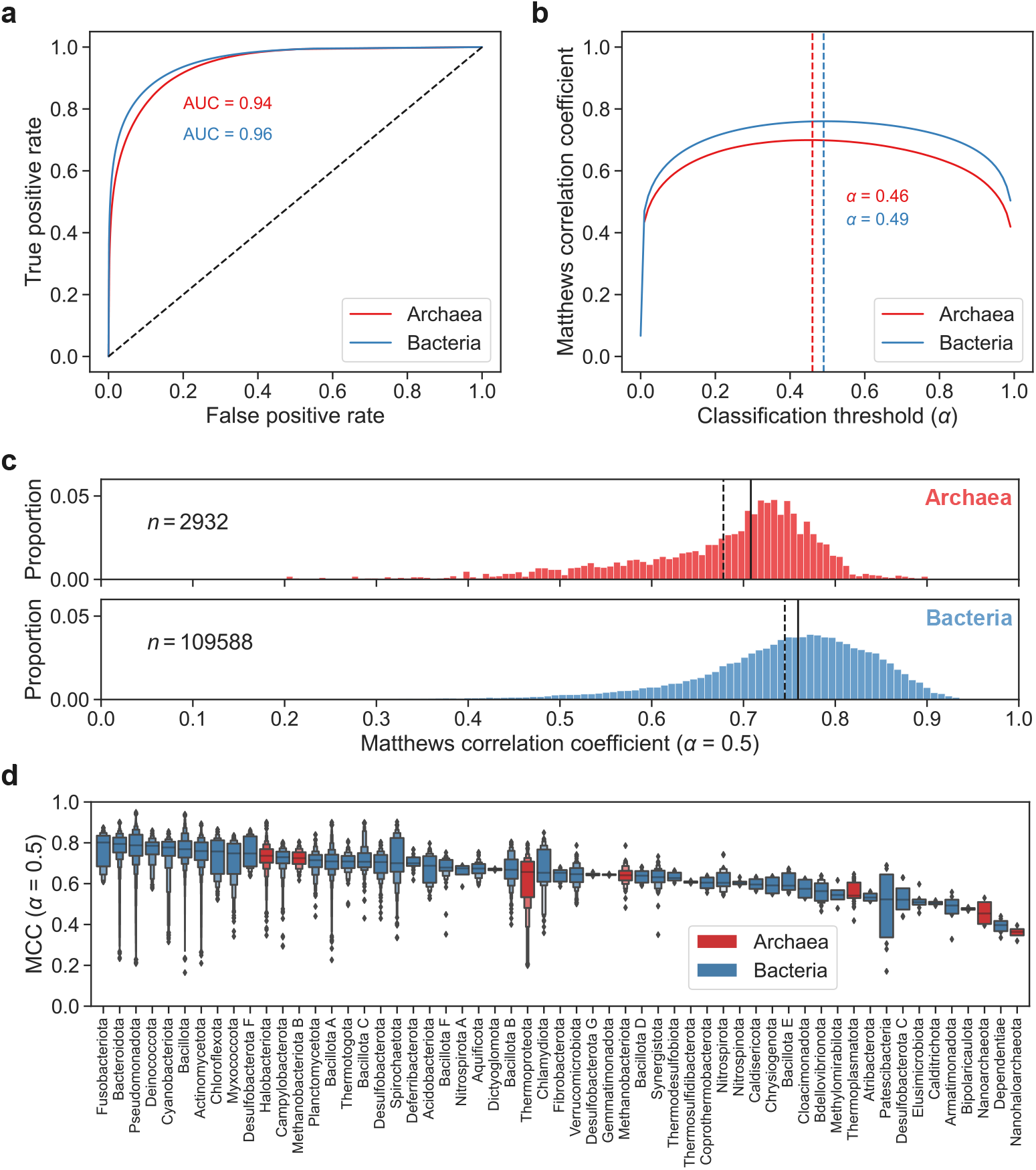
Classifying accurate archaeal and bacterial GEMs from ANN predictions. (a) ROC curves and AUCs for classification of GEMs by ANNs trained on MSAs from SINA and MAFFT. The dashed line indicates the performance of a random classifier. (b) MCC by classification threshold, *α*, for ANNs trained on MSAs from SINA and MAFFT. Dashed lines indicate optimal thresholds that maximize mean MCC. The curves shows the mean across sequence representations and test sets for archaea and bacteria. (c) MCCs for archaeal and bacterial GEMs classified by ANNs trained on MSAs from SINA and MAFFT with classification threshold *α* = 0.5. Solid and dashed lines indicated median MCC and mean MCC, respectively. (d) MCCs by phylum for archaeal and bacterial GEMs classified by ANNs trained on MSAs from SINA and MAFFT with classification threshold *α* = 0.5. Letter-Value plot showing median MCC with quantiles and outliers [25].

## Discussion

By extracting 16S rRNA gene sequences and GEMs from more than 15,000 reference and representative RefSeq genomes, we were able to train a simple ANN to predict microbial metabolism directly from a single gene sequence. This enables genome-scale simulation of microbes that lack genomic data but have been observed through 16S rRNA sequencing. Our trained ANNs can easily be applied to existing 16S sequencing data, and it is also easy and fast to retrain or fine-tune the ANNs on other data sets. This novel application of neural networks can be extended, e.g., to other marker genes, sequence representations, or genome-scale features beyond metabolism.

Reference and representative genomes from RefSeq [5] covered known archaeal and bacterial taxonomy well, and we were able to extract GEMs from all the genomes and 16S rRNA gene sequences from most of them. Thanks to CarveMe [3] and other automated metabolic reconstruction methods [26], building large collections of high-quality microbial GEMs from annotated genomes has become very fast. Our GEMs extend existing collections and can easily be reproduced or expanded by others [3]. For example, one could extend the training data substantially by building GEMs for larger collections of genomes, e.g., the full GTDB taxonomy [20], although adding genomes of lower quality could reduce the accuracy of ANN predictions. If one is specifically interested in high-quality predictions for the human gut microbiome, one could apply our approach to very large existing GEM collections of human gut microbes [8, 9].

ANNs trained on different sequence representations were similar in their predictions and accuracy, but unaligned sequences were outperformed by LLM embeddings, which in turn were outperformed by MSAs. It is notable that the network learned very similar representations of the relationship between 16S rRNA gene sequences and GEMs from unaligned sequences as well as from aligned but very different sequence representations. This indicates that the ANN was able to compensate for lack of alignment, at least to some extent, and that the network architecture, which enforces an increasingly sparse representation, helped make predictions quite robust to the input data. It is also notable that the ANN was able to extract more information from the embeddings of an LLM that was not trained specifically for this purpose than from the unaligned sequences, although still less than from the aligned sequences.

Classifying predictions from the most accurate trained ANNs, i.e., those using MSAs for training, produced ROC curves with very high AUC for both archaea and bacteria. This means that the true positive rate was very high compared to the false positive rate across reaction probability thresholds, i.e., that ANN predictions of reaction presence in a GEM were likely to be true. Here it is important to note that we never allowed any overlap of genomes or 16S rRNA gene sequences between training and test sets. MCCs were slightly lower than the AUCs because of the false negative rate, which was higher than the false positive rate across thresholds. This means that predictions of reaction absence were more likely to be wrong than predictions of reaction presence. Because of this, and to enable simulation of growth and metabolic fluxes with methods such as flux balance analysis [24], gap filling should be performed as a final step before application of GEMs [27].

There are of course several important limitations to our ANN-based approach for predicting GEMs from 16S rRNA gene sequences. First, it is completely dependent on large collections of high-quality genomes with corresponding models and marker gene sequences. Thus, our approach should not be viewed as a competitor to manual or automated genome-based approaches, but rather as a complement that can use existing high-quality data to extend the taxonomic scope of GEMs. Second, GEMs built manually or automatically from genomes are themselves limited by many sources of uncertainty [28]. This propagates to ANNs trained on GEMs and is likely amplified if genome quality is reduced. Finally, our approach has not been exhaustively optimized, meaning that there could be further room for improvement of predictions, e.g., by adjusting the network architecture or training parameters or data.

Despite these limitations, we have shown that, given sufficient high-quality training data, ANN-based prediction of GEMs from 16S rRNA gene sequences can be accurate and robust for archaea as well as bacteria. The importance of sufficient data is highlighted by the fact that the predictive accuracy of trained ANNs was significantly lower for archaea than for bacteria, although the predictive accuracy for archaeal GEMs was impressive given that only 4% of the genomes in the training data came from archaea. Future studies should aim to increase the taxonomic scope and quality of genomes and GEMs even further, which will require substantial experimental and computational efforts. Once sufficient genomic data are available for a group of organisms, the manual effort required for reconstruction can be reduced and perhaps eventually eliminated by training ANNs to predict genome-scale features for organisms lacking data. Notably, expanding the repertoire of high-quality GEMs for eukaryotes from the tens to the hundreds or thousands would open the door to genome-scale simulation of growth and metabolism for the whole tree of life, underscoring the importance of ongoing efforts to characterize eukaryotic diversity through genome sequencing [29].

## Methods

### Genomes, 16S rRNA gene sequences, and GEMs

We downloaded all 18,339 available archaeal and bacterial reference and representative genome assemblies from the NCBI RefSeq database [5] on September 4, 2023. To extract all rRNA gene sequences from the genomes, we first found all entries with feature type “rRNA” and “16S” in the attribute field of the GFF file of each genome and then extracted the DNA sequences of these entries from the corresponding nucleotide FASTA files. This produced 471,666 sequences that were filtered by length to eliminate any fragments or 5S and 23S subunits. Specifically, only the 26,893 entries (25,149 unique sequences) with lengths larger than 1,000 bp and smaller than 2,000 bp were kept. This was based on the observed distribution of rRNA sequence lengths and the fact that 1,500 bp is the approximate mean length of full 16S rRNA gene sequences [30]. We used CarveMe [3] with the Gurobi solver [31] and default settings to build a GEM capable of predicting growth and metabolic fluxes from each genome.

### Numeric representations of 16S rRNA gene sequences

We represented the 26,893 16S rRNA gene sequences left after filtering numerically in three different ways. First, we simply assigned an integer to each unique base or other symbol found in the sequences. We padded the resulting integer sequences with zeros at the end to give them all a length of 1,997, equal to the length of the longest sequence. Second, we computed embeddings for the sequences using GenSLM 25M [21], an LLM with 25 million parameters that is trained on a large and varied set of prokaryotic gene sequences. Embeddings were computed by taking the mean of the hidden states of the final ANN layer at the end of training, producing a floating-point vector with 512 elements. Finally, we aligned the sequences using two different multiple sequence alignment programs: SINA [22] and MAFFT [23]. We ran SINA through its command line interface using default settings and the SILVA 138.1 SSURef NR99 reference database [32], and we ran MAFFT through its web interface with strategy FFT-NS-2 and low-memory mode to allow alignment of a large number of sequences (https://mafft.cbrc.jp/alignment/server/large.html). We discarded any unaligned parts from the MSAs, leaving 11,246 aligned positions for SINA and 24,690 for MAFFT. We converted the MSAs to integer sequences as described above.

### ANN architecture and training

To predict GEMs from 16S rRNA gene sequences, we used an ANN architecture consisting of three fully connected linear layers and a final sigmoid layer. The input and output sizes of the ANN depend on the data used for training, specifically the chosen representation of gene sequences and the number of unique reactions present in GEMs, respectively. The first fully connected layer converts the input to 512 features and the second decreases this to 256 features. The third fully connected layer converts the 256 features to the output size, and the final sigmoid layer converts these weights to numbers between zero and one that can be interpreted as reaction probabilities. We trained this ANN multiple times on pairs of GEMs and 16S rRNA gene sequence representations. Specifically, we paired binary vectors indicating presence and absence of reactions in each of the 18,339 GEMs with (1) unaligned sequences, (2) LLM embeddings from GenSLM 25M, (3) MSA from SINA, and (4) MSA from MAFFT. GEMs without any mapped 16S rRNA gene sequences were filtered out, leaving 15,827 GEMs with 5,248 unique reactions that were paired with representations of the 25,149 unique 16S rRNA gene sequences. We randomly split the data into training and test sets ten times, preserving an approximate 9/1 size ratio while ensuring that none of the sequences or GEMs in the training set were also included in the test set. We trained the ANN using each training set and the Adam optimizer with default learning rate 10^−3^ until convergence of the binary cross-entropy training loss after 20 epochs. The ANN was implemented and trained in Python using CUDA through PyTorch [33].

### GEM prediction and evaluation

We used the trained ANNs to predict GEMs from the 16S rRNA gene sequences in the test set. Using 100 linearly spaced classification thresholds from zero to one, we classified each reaction as present or absent in each GEM if its predicted probability was higher or lower than the threshold, respectively. We compared the results of this classification to the true GEMs built from genomes and computed the number of true positives (TP), false positives (FP), true negatives (TN), and false negatives (FN). We produced a ROC curve by plotting TP rate against FP rate and computed AUC as well as MCC:

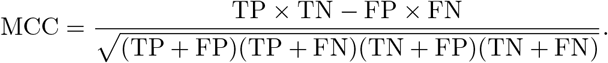

We found the classification thresholds that maximized MCC for archaea and bacteria and identified *α* = 0.5 as an approximately optimal and robust threshold.

### Computational resources

Analyses were performed on an AMD Ryzen Threadripper 5995WX CPU with 512 GB RAM and an NVIDIA RTX A4000 GPU with 16 GB VRAM. We used the GPU to compute LLM embeddings and train the ANN and the CPU for all other analyses.

## Data and code availability

Data and code that reproduce our results are available at gitlab.com/oveoyas/nnforge.

## Acknowledgements

We thank the rest of the BIAS and MEMO groups at NMBU for feedback and support. This work was funded by the Research Council of Norway grant 248792 (DigiSal) and the Novo Nordisk Foundation grant 0054575 (SuPA[cow]).

## Author contributions

O.Ø. and P.B.P. designed the study. O.Ø. developed the methods. O.Ø. and C.M.K. performed computational analyses. C.M.K. set up the computational resources. J.O.V. and P.B.P. secured funding. O.Ø., C.M.K., J.O.V., and P.B.P. wrote the manuscript.

